# An untargeted metabolomics approach to characterize dissolved organic matter in groundwater of the Samail Ophiolite

**DOI:** 10.1101/2022.11.09.515806

**Authors:** Lauren M. Seyler, Emily A. Kraus, Craig McLean, John R. Spear, Alexis S. Templeton, Matthew O. Schrenk

## Abstract

The process of serpentinization supports life on Earth and gives rise to the habitability of other worlds in our Solar System. While numerous studies have provided clues to the survival strategies of microbial communities in serpentinizing environments on the modern Earth, characterizing microbial activity in such environments remains challenging due to low biomass and extreme conditions. Here, we use an untargeted metabolomics approach to characterize dissolved organic matter in groundwater in the Samail Ophiolite, the largest and best characterized example of actively serpentinizing uplifted ocean crust and mantle. We found that dissolved organic matter composition is strongly correlated with both fluid type and microbial community composition, and that the fluids that were most influenced by serpentinization contained the greatest number of unique compounds, none of which could be identified using the current metabolite databases. Using metabolomics in conjunction with metagenomic data, we detected numerous products and intermediates of microbial metabolic processes and identified potential biosignatures of microbial activity, including pigments, porphyrins, quinones, fatty acids, and metabolites involved in methanogenesis. Metabolomics techniques like the ones used in this study may be used to further our understanding of life in serpentinizing environments, and aid in the identification of biosignatures that can be used to search for life in serpentinizing systems on other worlds.

## Introduction

Serpentinization is the geochemical process by which ultramafic rock reacts with water, resulting in the production of serpentine minerals and hydrogen gas (H_2_) (Sleep et al., 2004). High concentrations of H_2_ can further drive the reduction of dissolved inorganic carbon (DIC) to reduced carbon compounds including formate, methane, and carbon monoxide (McCollom and Seewald, 2001; Charlou et al., 2002; Proskurowski et al., 2008; McCollom, 2013.).

Chemolithotrophic microorganisms are able to use the products of these reactions as sources of carbon and reducing power (Schrenk et al., 2004; McCollom et al., 2007; Amend et al., 2011; Brazelton et al., 2011; Brazelton et al., 2012; Lang et al., 2012; Brazelton et al., 2013; Schrenk et al., 2013; Cardace et al., 2015). Serpentinization may be a widespread process in other locations in our solar system, including the subsurface of Mars, Enceladus, and Europa, making it a potential source of chemical disequilibria that may be harnessed by extraterrestrial life (Schulte et al., 2006; Vance et al., 2007; Klein et al., 2019; McCollom et al., 2022). Environments associated with serpentinization host microbial communities capable of a variety of metabolic strategies, including methanogenesis (Miller et al., 2016; Brazelton et al., 2017; Rempfert et al., 2017; Fones et al., 2019; Kraus et al., 2020), methane oxidation (Miller et al., 2016; Brazelton et al., 2017; Seyler et al., 2020a; Kraus et al., 2020), acetogenesis (Brazelton et al.; 2012; Rempfert et al., 2017; Suzuki et al., 2018; Fones et al., 2019; Seyler et al., 2020a; Coleman et al., 2022), and sulfate reduction (Sabuda et al., 2020; Nothaft et al., 2021; Templeton et al., 2021; Glombitza et al., 2021). However, the highly alkaline conditions resulting from the serpentinization reaction, coupled with the precipitation of available DIC to calcite at high pH (Barnes et al., 1978), creates a challenging environment for microbial life (Schrenk et al., 2013). The physiological strategies used by microbial communities to overcome these challenges are not well understood.

Metabolomics is a rapidly advancing tool in the field of microbial ecology (van Santen et al., 2020; Bauermeister et al., 2021; Oyedeji et al., 2021). The detection and annotation of metabolites using mass spectrometry has allowed researchers to demonstrate the existence of complete metabolic pathways not commonly detected by genome annotation (Tang et al., 2009), to describe the function of theoretical pathways (Peyraud et al., 2009), and to discover entirely novel pathways (Fürch et al., 2009; Liu et al., 2016). When integrated with other -omics approaches, including metagenomics and metatranscriptomics, metabolomics is a powerful tool for describing metabolic activity in microbial communities (Turnbaugh and Gordon, 2008; Beale et al., 2016; Porcar et al., 2018). Metabolomic profiling of microbial communities in extreme (e.g., hyperalkaline) environments can assist in understanding the metabolic pathways used by microbial communities to adapt to extreme conditions, and the mechanisms and degree to which metabolites may be preserved under these conditions (Blanchowitz et al., 2019). Metabolomics thus holds great potential in the search for biomarkers with astrobiological applications (Seyler et al., 2020b).

In this study, we used metabolomics to characterize dissolved organic matter derived from microbial metabolic activity in the Samail Ophiolite groundwaters in the Sultanate of Oman, which is the largest and best exposed ophiolite in the world. The Samail Ophiolite consists of a large (~15,000 km^3^) block of ocean crust and upper mantle (Nicolas et al, 2000) that was rapidly emplaced onto the Arabian continental plate approximately 70 million years ago (Glennie et al., 1973; Coleman, 1981; Tilton et al., 1981; Hacker et al., 1996). Here, mafic and ultramafic rocks that once underwent alteration in seafloor hydrothermal systems continue to be altered by active serpentinization on the continent (Canovas et al., 2017; Keleman et al., 2021). We used an untargeted metabolomics approach (Patti et al., 2012) to characterize compounds in the dissolved organic matter (DOM) pool of groundwaters collected from varied geochemical conditions, using metagenomic data acquired from biomass filtered from the same samples to verify the annotation of metabolomic features. To our knowledge, this is the first such study attempting to describe biomolecules that make up the DOM pool in the Samail Ophiolite groundwater.

## Methods

### Site Description

The Samail Ophiolite provides a complete cross-section through 7 km of oceanic crust and 15 km of underlying upper mantle rocks (Boudier and Coleman, 1981; Coleman and Hopson, 1981; Lippard et al., 1986; Glennie, 1996; Nicolas et al., 2000). Partially serpentinized peridotite in the ophiolite actively undergoes hydration and carbonation reactions at low temperatures (≤60°C), producing hyperalkaline fluids enriched in dissolved methane and hydrogen gas in the subsurface (Barnes et al., 1978; Neal and Stanger, 1983, 1985; Clark and Fontes, 1990; Kelemen and Matter, 2008; Kelemen et al., 2011, 2013; Paukert et al., 2012; Streit et al., 2012; Miller et al., 2016). The ophiolite’s size, exposure, and accessibility create unique opportunities for studying a broad range of geologic and hydrologic conditions supporting subsurface microbial life (Rempfert et al., 2017). Previous geochemical studies of the Samail Ophiolite groundwater observed a strong partitioning between two subgroups of peridotite-hosted fluids and classified them as “Type I/alkaline” (pH 8-10) or “Type II/hyperalkaline” (pH > 10), while wells positioned at the “contact” between peridotite and gabbro displayed more geochemical variability (Rempfert et al., 2017). Alkaline waters have greater availability of oxidants and DIC and more meteoric influences than hyperalkaline, while contact waters appear to be a mixture of the two (Nothaft et al., 2021). We continue to use these designations to refer to well types throughout this study.

### Sampling

In February 2017, a submersible pump was used to collect water samples from five previously drilled wells in the Samail Ophiolite (Table 1). Approximately 100 L of flush water were pumped from at least 20 meters below the air-water interface in each well prior to sample collection. Sample volume varied according to the amount of water that could be acquired from each well while continuously pumping: 20 L were collected from NSHQ14, 7 L from WAB71, 18 L from WAB55, 4 L from WAB104, and 20 L from WAB105. Each sample was filtered through 0.3 μm glass fiber filters in a muffled stainless steel inline filter holder using PTFE tubing. Approximately 30 ml filtrate was quenched and stored in liquid nitrogen for transport back to the laboratory, where total organic carbon analysis was performed on a Shimadzu total organic carbon analyzer. The rest of the filtrate was acidified to pH 3 using concentrated HCl for solid phase extraction in the field. Biomass for DNA extraction was collected using in-line 0.2 μm polycarbonate filters as described previously (Fones et al., 2019). 10 mL samples of unfiltered waters from each well were preserved in the field for enumeration of planktonic cells by adding formaldehyde to a final concentration of 10% vol/vol, and transported back to the laboratory for direct cell counting via 4ʹ,6-diamidino-2-phenylindole (DAPI) staining on black polycarbonate 0.2 μm filters. Water temperature, conductivity, pH, oxidation-reduction potential (ORP), and dissolved oxygen concentration were measured during sampling using a Hach HQ40D Portable Multi Meter (Fones et al., 2019).

**Table 1.** Geochemical data of wells sampled for metabolomics.

| site | lithology | pump<br>depth<br>(m) | pH | T<br>(°C) | Cond<br>(μS/<br>cm) | ORP<br>(mV) | Cell Counts<br>(per ml) | TOC<br>(mg/L) | SPE<br>Volume<br>(L) | TOC of<br>sample<br>(mg) |
| --- | --- | --- | --- | --- | --- | --- | --- | --- | --- | --- |
| NSHQ14 | peridotite | 50 | 11.1 | 34.4 | 493 | -415 | 3.04E+05 | 1.4470 | 20 | 28.94 |
| WAB71 | peridotite | 50-70 | 10.6 | 34.5 | 1803 | -86 | 2.79E+05 | 2.5130 | 7 | 17.591 |
| WAB55 | contact | 30 | 9.2 | 34.7 | 1171 | 110 | 6.79E+04 | 0.3560 | 18 | 6.408 |
| WAB104 | peridotite | 70 | 8.5 | 33.4 | 493 | 180 | 2.38E+05 | 0.6146 | 20 | 12.292 |
| WAB105 | peridotite | 50 | 8.3 | 31.6 | 448 | 178 | 1.80E+05 | 0.5756 | 4 | 2.3024 |

### Metabolomics

Dissolved organics were captured from the acidified filtrate on a solid phase extraction Bond Elut-PPL cartridge (Agilent) (Dittmar et al., 2008; Fiore et al., 2015; Seyler et al., 2020a), using 1/8” × 1/4” PTFE tubing to pull the supernatant into the cartridge to minimize the possibility of contamination from plastic leaching, and Viton tubing to remove the discarded flow-through via peristaltic pump (flow rate not exceeding 40 ml/min). The cartridges were then rinsed with at least 2 cartridge volumes of 0.01 M HCl to remove salt, and the sorbent dried with air for 5 minutes. Dissolved organic matter (DOM) was eluted with 2 ml 100% methanol via gravity flow into muffled amber glass vials. These extracts were then shipped back to the laboratory at Michigan State University, where they were dried down via vacuum centrifugation, and resuspended in 495 ul 95:5 water:acetonitrile and 5 ul 5 μg/ml biotin-(ring-6,6-d_2_) standard (Rabinowitz and Kimball, 2007; Fiore et al., 2015). This protocol was repeated for 4 L milliQ water as an extraction blank. In addition, an extraction solvent blank was run consisting of 495 μl of 95:5 water:acetonitrile plus 5 μl 5 μg/ml biotin-(ring-6,6-d_2_).

Samples were analyzed via quadrupole time-of-flight liquid chromatography tandem mass spectrometry (QToF-LC/MS/MS) on a Waters Xevo G2-XS UPLC/MS/MS instrument at the MSU Metabolomics Core facility. Each sample was separated chromatographically in an Acquity UPLC BEH C18 column (1.7 μm, 2.1 mm × 50 mm) using a polar/non-polar gradient made up of 10 mM TBA and 15 mM acetic acid in 97:3 water:methanol (solvent A), and 100% methanol (solvent B). The gradient was run at 99.9/0.1 (% A/B) to 1.0/99.0 (% A/B) over 9 min, held an additional 3 min at 1.0/99.0 (% A/B), then reversed to 99.9/0.1 (% A/B) and held another 3 min. At a rate of 400μL/min, the sample was fed into a quadrupole time-of-flight mass spectrometer using a Waters Xevo G2-XS MS/MS in negative ion mode using a data independent collection method (m/z acquisition range 50 to 1500 Da). Mass spec data were then imported into Progenesis QI software for statistical analysis and correlation. Quality control was performed by checking ion intensity maps and producing alignments of all sample runs. An aggregate data set was then created from the aligned runs to produce a map of compound ions that was applied to each sample for peak matching. Compound ions were deconvoluted and ion abundance measurements normalized so that runs could be compared. This approach yielded a table of peak areas that are identified by a unique mass to charge (m/z) and retention time (rt) pair referred to as features. Multivariate statistics was then performed on the data including principal components analysis and correlation analysis. A table of relative abundances of features was generated for further analysis. Features found in the blanks were subtracted from the sample data (Broadhurst et al., 2018). In order to be considered present in a sample, a feature had to have at least a two-fold greater abundance in the sample than in both blanks. Features were putatively annotated in Progenesis QI against the ChemSpider database using mass, charge, retention time, and fragmentation spectra. Further annotation of features was achieved using mummichog (Li et al., 2013), which predicts functional activity by compiling metabolic networks using mass spec data. Whenever possible, we used autonomous METLIN-Guided In-source Fragment Annotation (MISA) (Domingo-Almenara et al., 2019), an algorithm that compares detected features against low-energy fragments from MS/MS spectra in the METLIN database (Smith et al., 2005), to increase our confidence in the putative annotation of representative metabolites; however, most of the putative metabolites did not have MS/MS spectral data available in the database. Raw spectrometry data are available in the MetaboLights database (Haug et al., 2020) under study identifier MTBLS6081.

### Metagenome Binning

DNA extraction, generation of sequencing libraries, and analysis of metagenomic sequence data, including binning, was described previously (Fones et al., 2019; Kraus et al., 2020).

Bins were taxonomically classified using GTDB-Tk (Chaumeil et al., 2019). Bin quality was assessed using CheckM version 1.0.11 (Parks et al., 2014); bins with 50% or greater completion were selected for further downstream analyses. The relative abundance of each bin in the metagenomes was calculated using a competitive recruitment approach. A bowtie2 index was built from a concatenated file of all bin contigs using bowtie2-build (Langmead and Salzberg, 2012). Quality-filtered metagenome reads were mapped against the bin bowtie2 index using bowtie2 with the–no-unal flag. The resulting SAM files (containing only the reads which mapped) were converted to BAM files, then BamM (https://github.com/Ecogenomics/BamM) was used to remove all alignments with <95% identity and <75% alignment coverage. The number of reads in each remaining BAM file was counted using Binsanity-profile (v0.3.3) (Graham et al., 2017). The read counts were then used to calculate the normalized relative fraction of the metagenomes that mapped to the bins using the following equation:

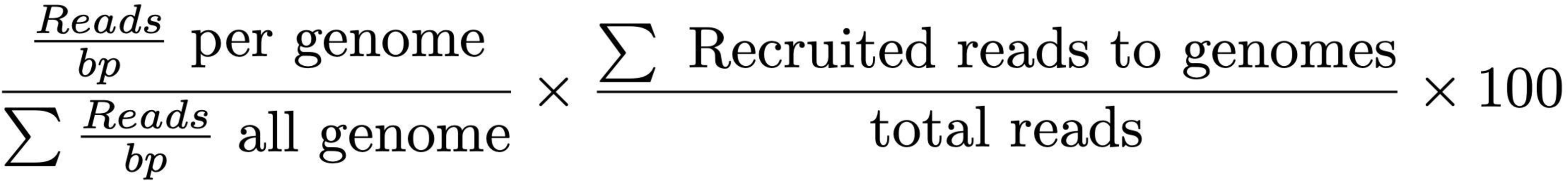

Z-scores of the normalized relative fractions were calculated using the average and sample standard deviation of the relative fraction of each bin across all metagenomes. The bin relative abundance data was run through the Nonnegative Matrix Factorization (NMF) package (Gaujoux and Seoighe, 2010) in R to run a standard nonnegative matrix factorization algorithm based on the Kullback-Leibler divergence (Brunet et al., 2004). This algorithm generated a heatmap of the bins across samples and a dendrogram using a Pearson correlation distance function. Bins that were representative of particular wells or geochemical conditions were selected using the method defined by Carmona-Saez et al. (2006). MetaSanity (Neely et al., 2019) was used to annotate functional orthologies against the KEGG database (Kanehisa et al., 2019); these annotations were run through KEGG-Decoder (Graham et al., 2018) to determine the completeness of metabolic pathways of interest. Genes and pathways not detected by KEGG-Decoder were searched for in the bins using protein sequences available in UniProt (The UniProt Consortium, 2020) and the Conserved Domain Database (Lu et al., 2020), the MUSCLE alignment tool (Edgar, 2004), and the hmmsearch function in HMMER v 3.3.2 (Eddy, 2008).

### Cross-referencing Metabolomic and Metagenomic Data

Metabolomic feature abundance and bin abundance were standardized for ordination according to Legendre and Gallagher (2001) using the decostand function in the R vegan package (v2.4-2; https://www.rdocumentation.org/packages/vegan). Regularized Discriminant Analysis (RDA) was performed on the standardized data using the klaR package (v0.6-14; https://www.rdocumentation.org/packages/klaR). These RDAs were then subjected to Procrustes rotation using function Procrustes with symmetric scaling.

Feature annotations acquired using mummichog were cross-referenced with intermediate compounds from pathways identified in representative bins. As for the bins, NMF analysis was performed, and a heatmap of features across samples and a dendrogram using a Pearson correlation distance function were generated. Features that were representative of particular wells or geochemical conditions were extracted using the procedure described by Kim and Park (2007).

## Results

Total organic carbon (TOC) was highest in WAB71 (2.5130 mg/L), followed by NSHQ14 (1.4470 mg/L), WAB104 (0.6146 mg/L), WAB105 (0.5756 mg/L), and WAB55 (0.3560 mg/L) (Table 1). A total of 2483 features were detected in the mass spectrometry data across all wells (Supplementary Table 1). Principal coordinate analysis of the mass spectrometry data grouped WAB71, WAB55, WAB104, and WAB105 together and set NSHQ14 apart from the other wells (Figure 1). A Pearson correlation of feature abundance across wells followed by NMF analysis of feature abundance using the standard algorithm (Brunet et al., 2004) and a factorization rank of three clustered the wells into three “levels”: NSHQ14, WAB71, and the alkaline/contact wells (WAB104, WAB105, WAB55) (Figure 2). A factorization rank of two resulted in WAB71 being clustered with WAB104, WAB105, and WAB55 (data not shown). NMF analysis also identified features which were representative of (i.e., most abundant in and most likely to be found in) each level. In our analyses, 48 features were designated as representative of NSHQ14, only one of which could be annotated by mummichog; two features were representative of WAB71, neither of which could be annotated by either Progenesis QI or mummichog; and the remaining wells had 90 representative features, eight of which could be annotated by mummichog (Table 2). We also searched the mass spectrometry data for features which were only detected in one well (unique features). NSHQ14 had the greatest number (200) of unique features, WAB71 had 49, WAB104 had 15, WAB105 had five, and WAB55 had no unique features; none of these unique features could be annotated by either Progenesis QI or mummichog (data not shown).

**Figure 1.**
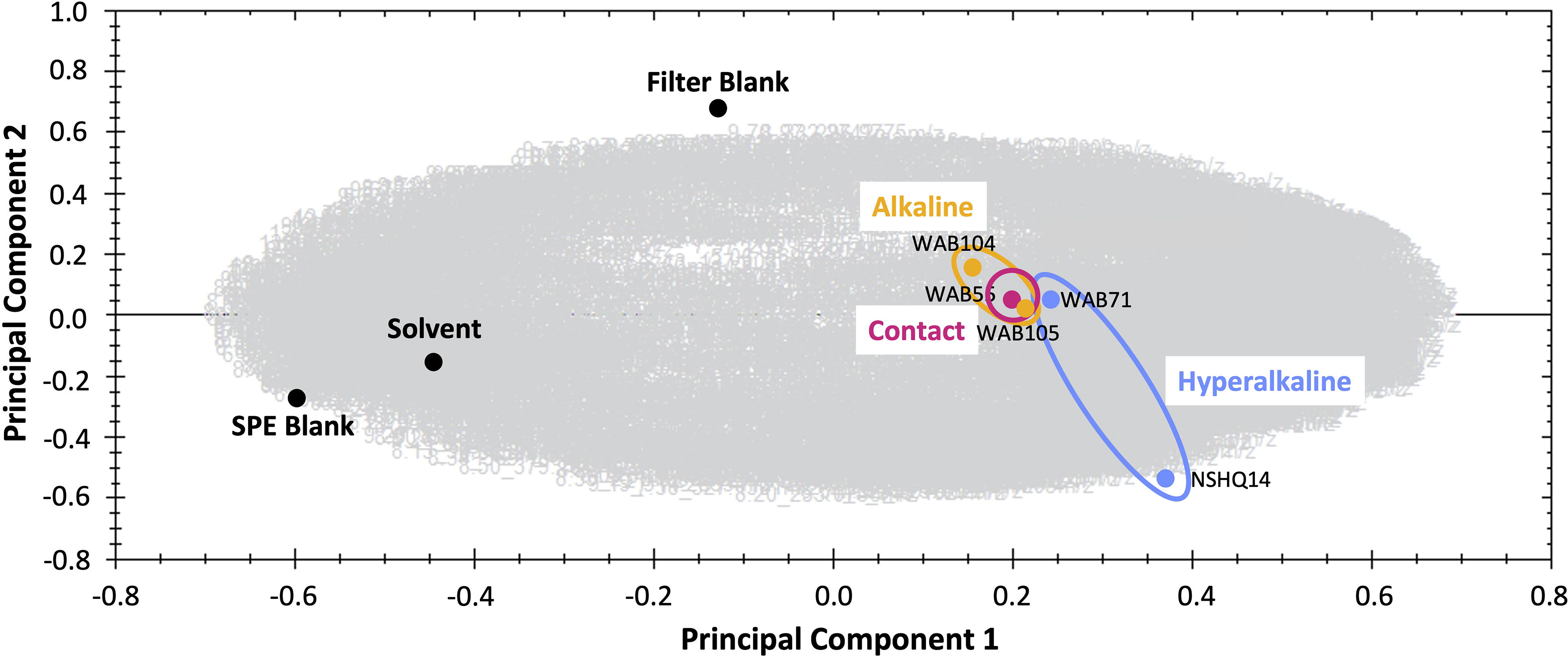
Principal component analysis comparing metabolomes of five sampled wells and blanks. The gray area is a cloud of all metabolomic features across all samples.

**Figure 2.**
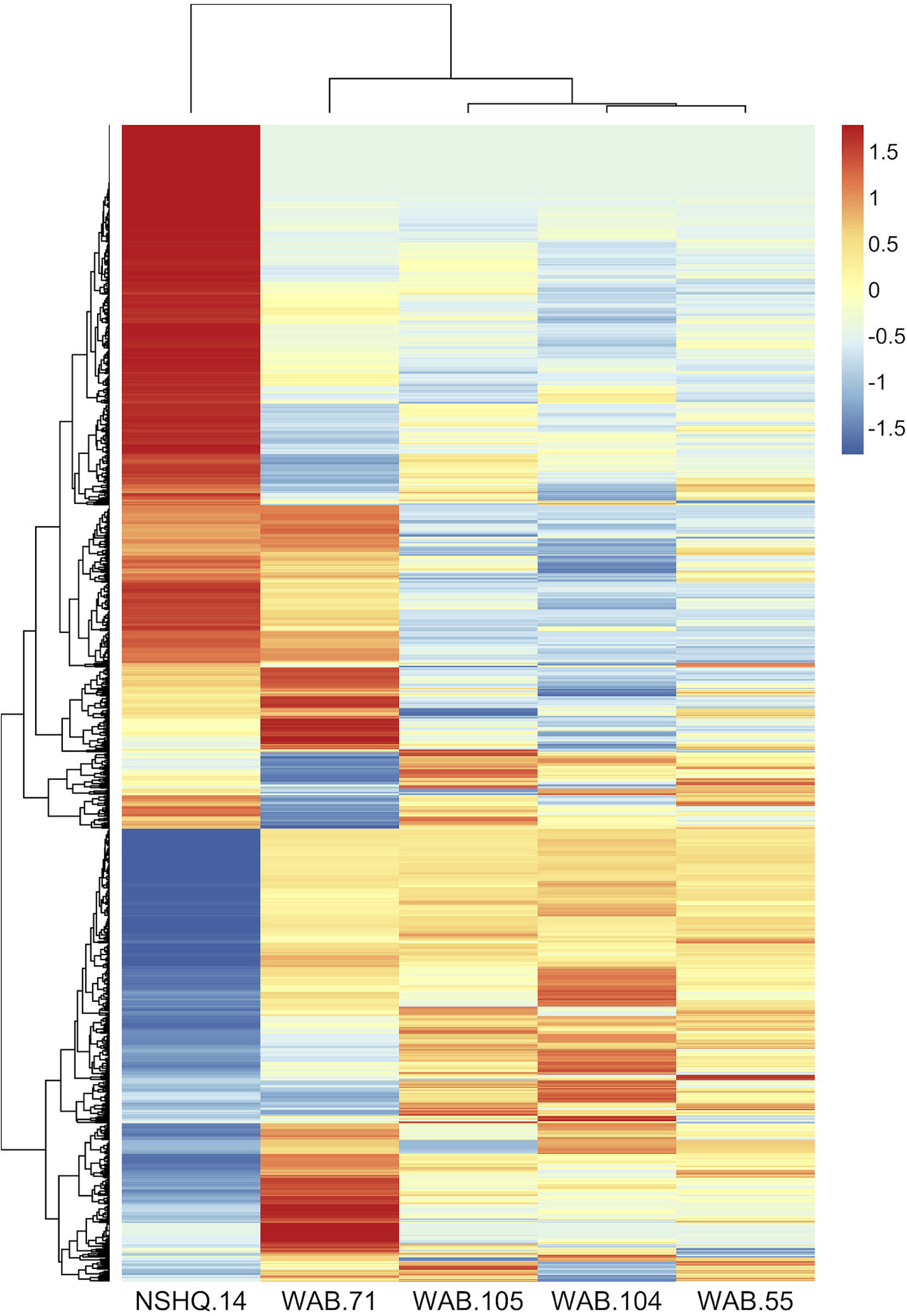
Z-scores of metabolomic feature abundance across wells clustered using a Pearson correlation, indicate peaks in the mass spectrometry data are differentially more abundant in some wells than in others. In particular, NSHQ14 had many unique features, most of which could not be identified.

In contrast to the metabolomic features, metagenomic bins tended to cluster by well (Figure 3). NMF analysis clustered the bins according to fluid type, and identified bins which were representative of (i.e., most abundant in and most likely to be found in) the hyperalkaline (NSHQ14 and WAB71), alkaline (WAB104 and WAB105), or contact (WAB55) wells. Superimposing the ordination plots of feature abundance and bin abundance data using Procrustes analysis showed that variance in the DOM pool could largely be attributed to variance in bin abundance and that the wells clustered according to fluid type, with the exception of WAB71, whose DOM pool clustered with the alkaline wells (Figure 4). Procrustes analysis also distinguished the DOM pool of WAB55 from those of NSHQ14 and WAB71, WAB 104, and WAB105 (Figure 4).

**Figure 3.**
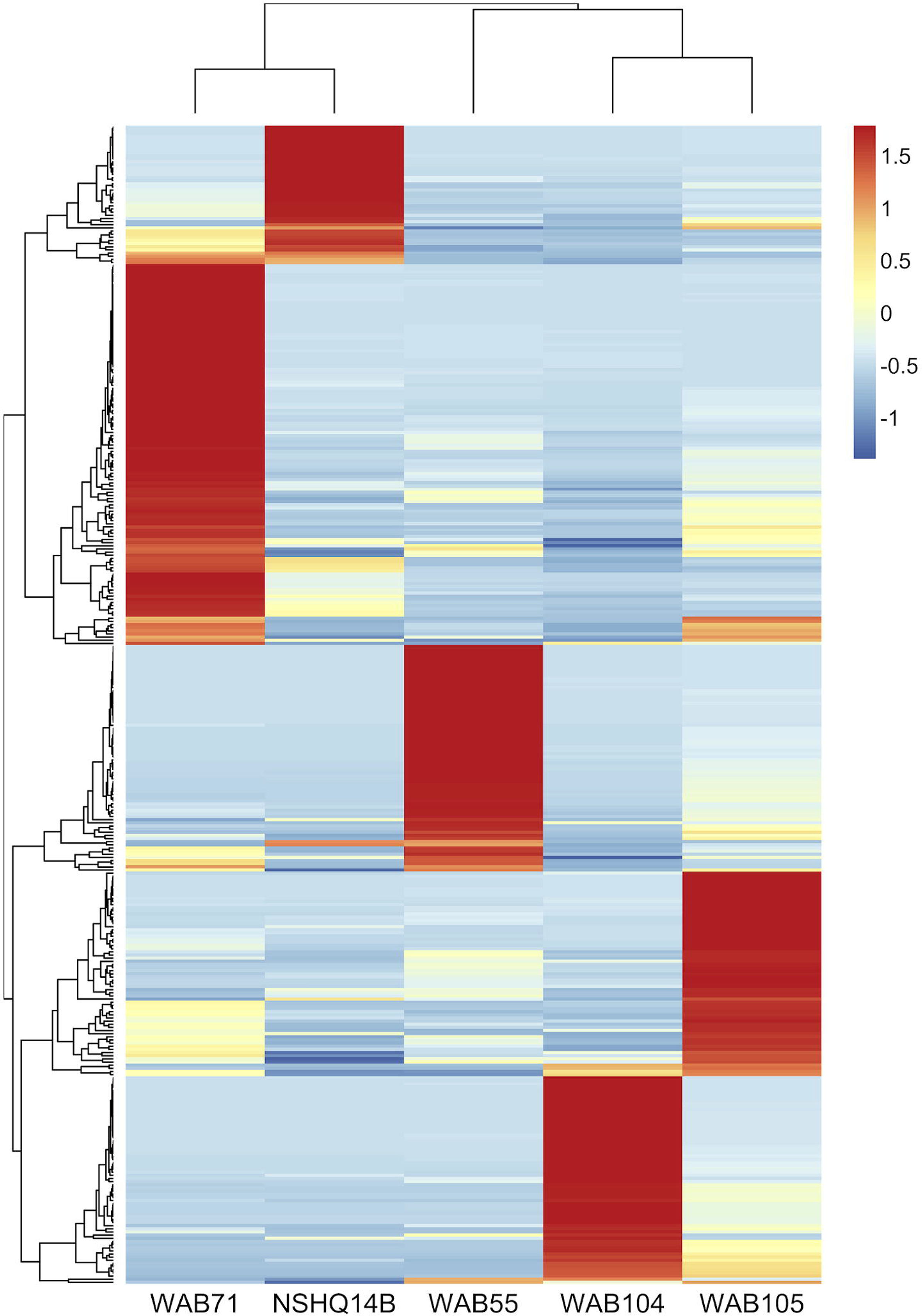
Z-scores of bin abundance across wells, clustered using a Pearson correlation. Wells clustered according to fluid type: hyperalkaline (NSHQ14 and WAB71), alkaline (WAB104 and WAB105) and contact (WAB55). WAB71 had the greatest number of bins which were more abundant in this well than any other well, while NSHQ14 had the fewest.

**Figure 4.**
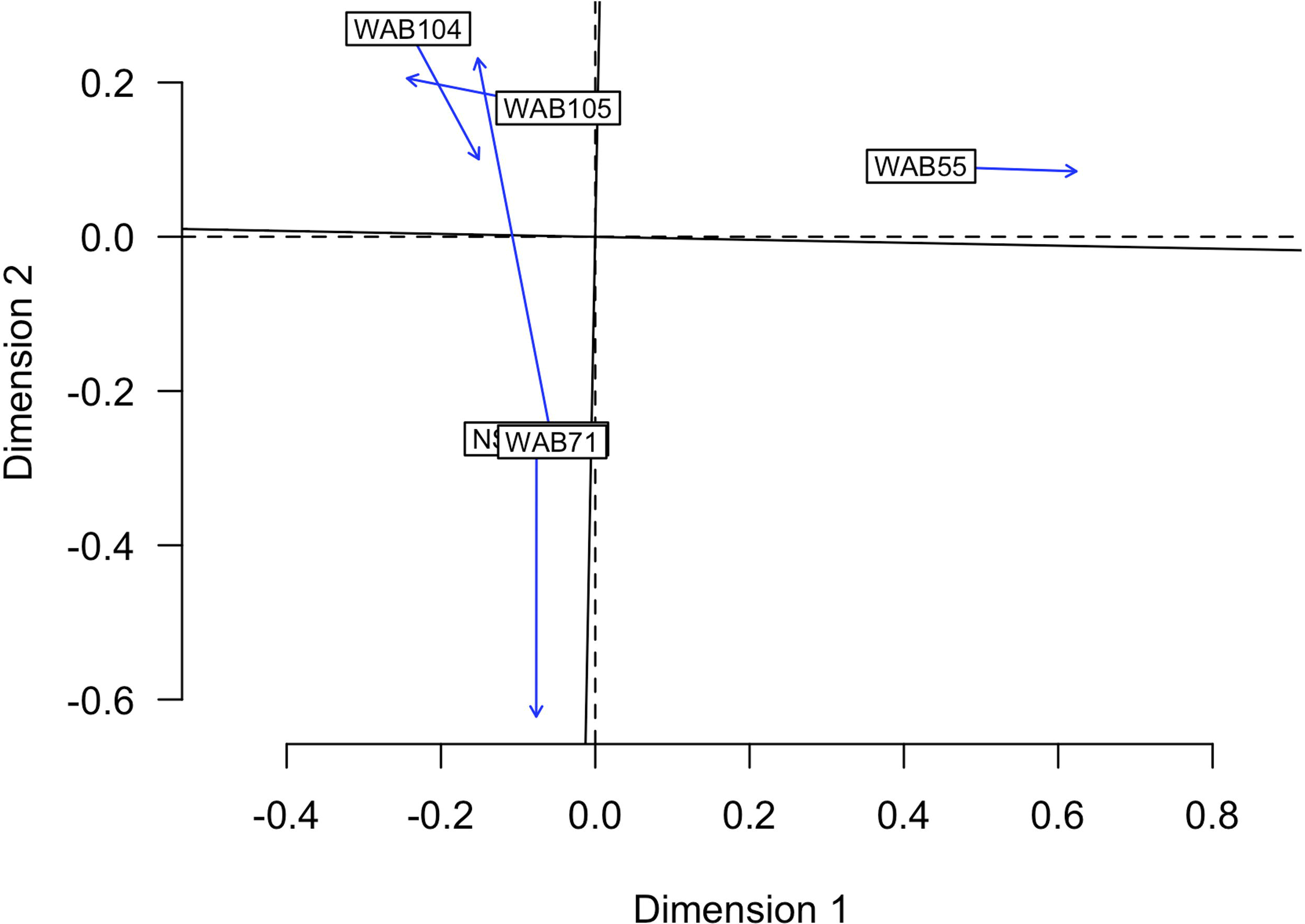
Procrustes analysis in which the metabolomic feature abundance and bin abundance ordinations are optimally superimposed through rotating and rescaling, showing that metabolomic feature abundance and bin abundance are correlated. Hyperalkaline wells and alkaline wells cluster together, while the contact well is distinct from both. Labeled boxes indicate the position of the samples in the first ordination, and arrows point to their positions in the target ordination.

1625 of the 2483 features in the mass spectrometry data were putatively annotated by Progenesis QI, by matching feature mass to compounds in the ChemSpider database. After removing features that appeared in the extraction blanks, the list of features was reduced to 953, 206 of which could be annotated by Progenesis QI. In contrast, 119 of these 953 features were able to be annotated by mummichog, which annotates metabolomic data by using network analysis and metabolite prediction to resolve ambiguity in metabolite annotation. Because Progenesis QI relies on mass matches to the ChemSpider database, which includes synthetic and theoretical compounds in addition to naturally-occurring metabolites, we primarily relied on the mummichog annotations for our analyses. We cross-referenced metabolic pathways annotated in the representative bins by KEGGDecoder with the features that could be annotated by mummichog to select features in the data on which to further focus our analyses. We then used the MetaCyc database (Caspi et al., 2014) to identify specific reaction pathways involving these metabolites and confirmed the presence of the relevant genes in the annotated MAGs. Pathways of interest and their annotated features in the mass spectrometry data are outlined below.

### Carbohydrates and Organic Acids

Putative annotations of mass spectrometry features included glycan sugars (L-fuculose, L-rhamnose, L-rhamnulose) and their degradation products (L-fucono-1,5-lactone and 2-dehydro-3-deoxy-L-rhamnonate), sugar alcohols, polysaccharides, butanoate, and isocitrate (Figure 5). We further confirmed the annotation of isocitrate using MISA.

**Figure 5.**
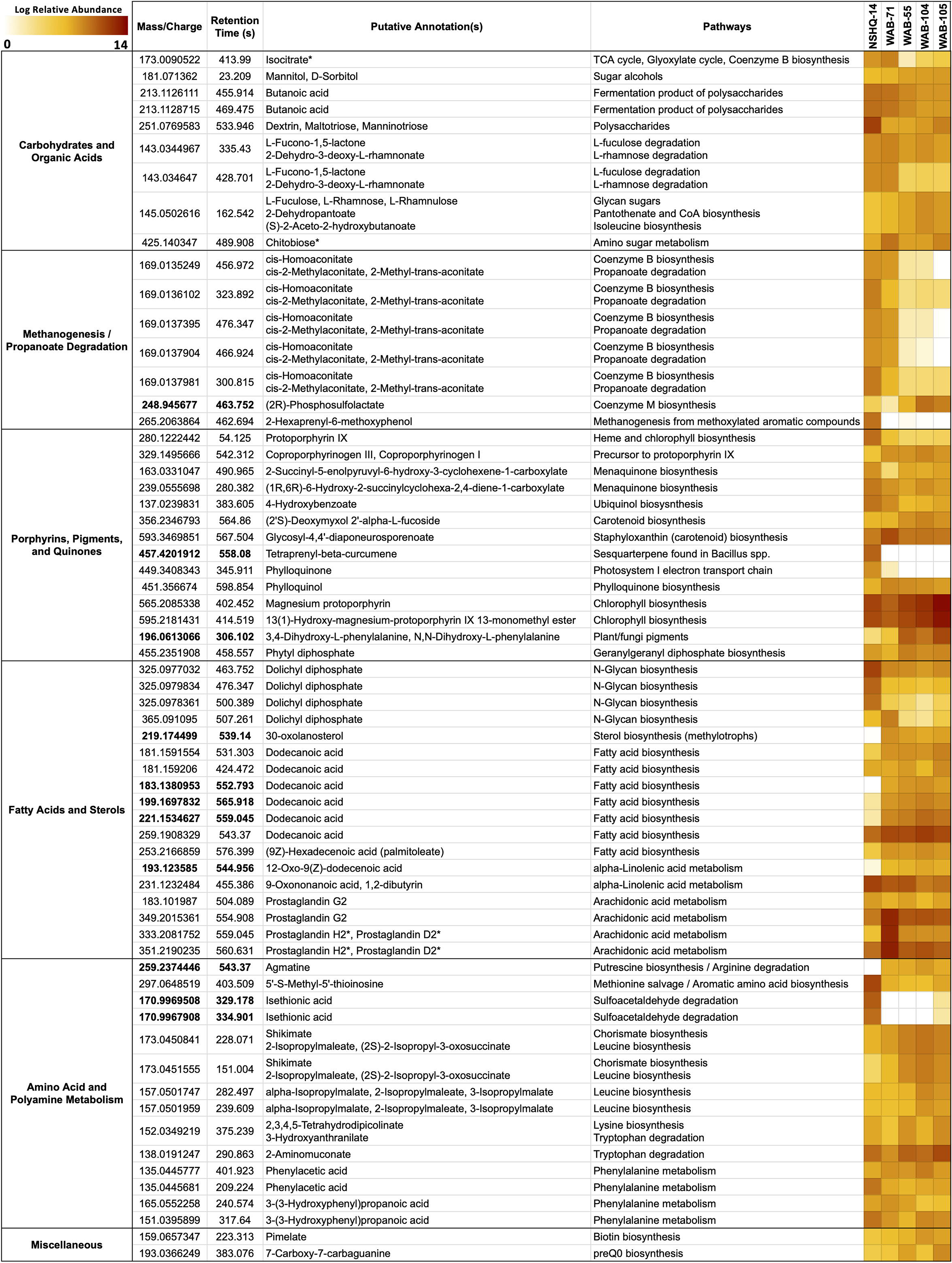
Features cross-referenced between KEGG results and mummichog annotations. Features are grouped by putative annotation/function. Diagnostic features appear in bold. Feature annotations confirmed by MISA are indicated with an asterisk (*). Feature abundance is expressed as the log of the normalized relative abundance.

Isocitrate was more abundant in the hyperalkaline wells than in the contact or alkaline wells. Isocitrate is an intermediate of multiple metabolic pathways, most notably the citric acid (TCA) cycle, the reverse TCA (rTCA) carbon fixation pathway, the glyoxylate cycle, and coenzyme B biosynthesis in methanogens. A partial TCA cycle was annotated in nearly every bin generated from the metagenomic data, but the rTCA pathway was not identified in any of the bins (Figure 6). The glyoxylate cycle, which uses five of the eight enzymes of the TCA cycle to synthesize macromolecules from two-carbon compounds (e.g., acetate), was annotated in 25 bins, three of which were representative bins of the alkaline wells, three which were representative of the contact well, and one which was representative of the hyperalkaline wells (Figure 6). A complete pathway for coenzyme B biosynthesis was detected in only one bin, a *Methanobacterium* that was representative of NSHQ14 (Figure 6).

**Figure 6.**
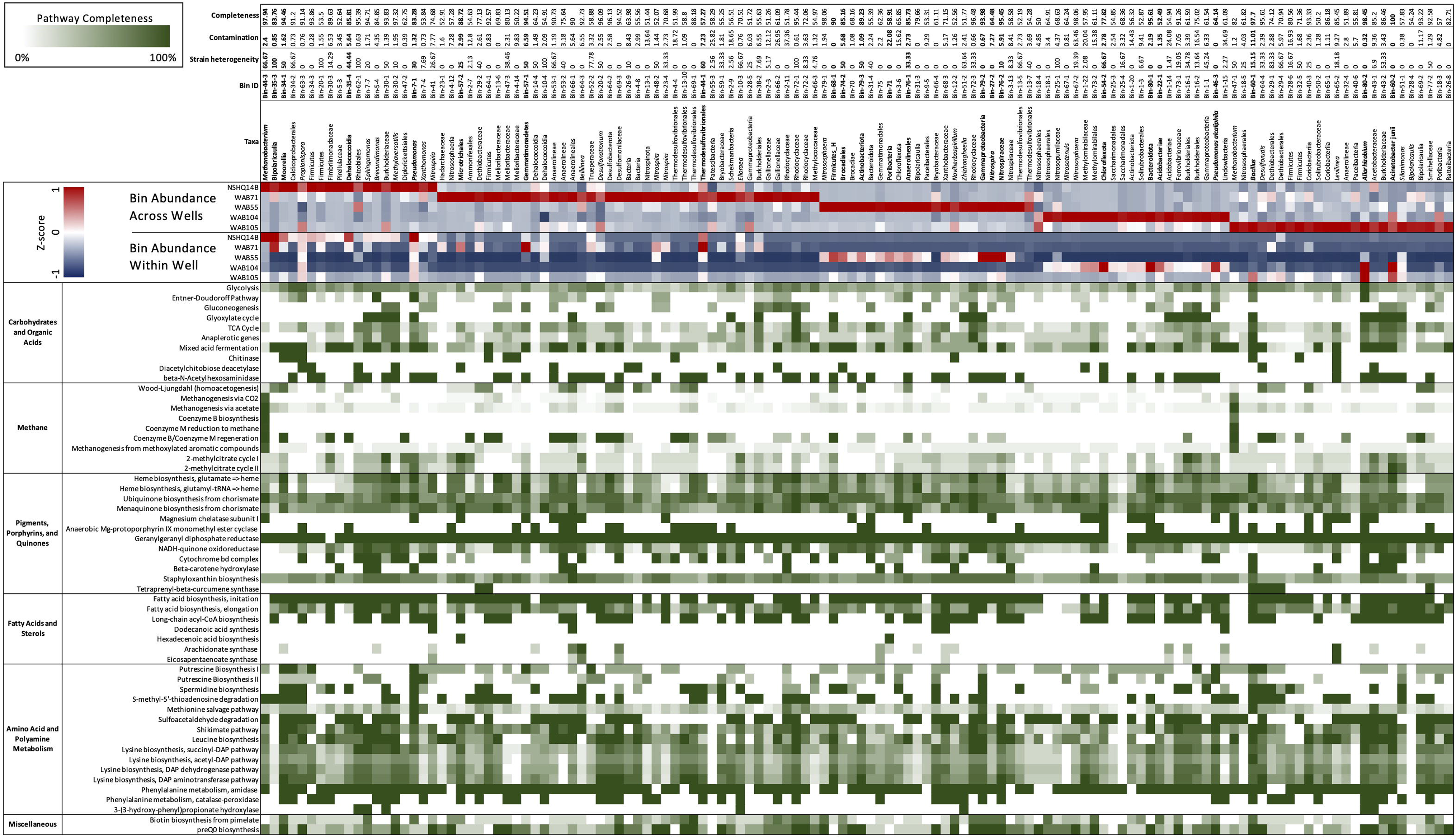
Metabolic genes and pathways of interest, annotated in each bin using KEGGDecoder and HMMER. Percentage completion of each pathway is indicated in shades of green. Z-scores of bin abundance within each well and across all wells are indicated in a blue/red heatmap at the top of the figure (negative z-scores in blue, positive z-scores in red).

Butanoate was also most abundant in the hyperalkaline wells. Butanoate may be produced by the fermentation of pyruvate, but while fermentation pathways were found in 31 (23 representative) bins (Figure 6), the complete set of enzymes required for pyruvate fermentation to butanoate were not detected in any of the bins (Supplementary Table 3).

A complete pathway for L-fuculose or L-rhamnose degradation was not annotated in any of the bins (Supplementary Table 3). The gene for D-threo-aldose 1-dehydrogenase, which produces L-fucono-1,5-lactone during L-fucose degradation, was only annotated in three bins, but the gene for L-fuconolactonase, which further degrades L-fucono-1,5-lactone to L-fuconate, was annotated in fifteen bins (Supplementary Table 3). Similarly, the gene for the L-rhamnonate dehydratase enzyme that produces 2-dehydro-3-deoxy-L-rhamnonate was only annotated in two bins, but the enzyme that acts on this product, 2-dehydro-3-deoxy-L-rhamnonate dehydrogenase, was annotated in seven bins (Supplementary Table 3).

Chitobiose, a disaccharide produced by the hydrolysis of chitin, was also annotated in the mass spectrometry data (Figure 5). MISA further confirmed the annotation of chitobiose; however, we question the validity of the annotation as the peaks eluted very far apart, suggesting that they did not arise from the same parent molecule (Supplementary Figure 1). Genes for chitinase and diacetylchitobiose deacetylase, enzymes that hydrolyze chitin and chitobiose, were identified in eleven and sixteen bins respectively, though only two bins contained both (Figure 6). Beta-*N*-acetylhexosaminidase, an exoglycosidase that catalyzes the hydrolysis of chitobiose, was detected in 54 out of 128 total bins, 14 of which were representative bins (6 of the 7 alkaline representative bins, 3 hyperalkaline representative bins, and 5 contact representative bins) (Figure 6). Genes for enzymes involved in the catabolism of other polysaccharides, including alpha-amylase, glucoamylase, beta-glucosidase, D-galacturonate epimerase, D-galacturonate isomerase, oligogalacturonide lyase, and pullulanase were also detected in the metagenomic data, and tended to be associated with bins that displayed the greatest abundance in the hyperalkaline wells (Supplementary Table 3).

### Methanogenesis/Propanoate Degradation

Five features were putatively annotated as either cis-homoaconitate, an intermediate in coenzyme B biosynthesis, or cis-2-methylaconitate/2-methyl-trans-aconitate, intermediates in propanoate degradation to pyruvate and succinate via the 2-methylcitrate cycle (Figure 5). These features were more abundant (by up to eight orders of magnitude) in the hyperalkaline wells compared to the contact and alkaline wells. An intermediate of coenzyme M production, (2R)-phosphosulfolactate, was also putatively annotated in the mass spectrometry data and was detected in all of the wells, though it was most abundant in the alkaline wells and was designated a representative feature of the alkaline/contact wells by NMF analysis (Figure 5). A feature putatively annotated as 2-hexaprenyl-6-methoxyphenol, an intermediate in methanogenesis from methoxylated aromatic compounds, was almost exclusively found in NSHQ14 (Figure 5). The enzyme that produces this intermediate has not been characterized.

Complete pathways for methanogenesis via CO_2_ and coenzyme M reduction to methane were found in two out of the 128 bins analyzed (Figure 6; Fones et al., 2019). Both of these bins were taxonomically annotated as Euryarchaeaota belonging to the Methanobacteriales; one bin was designated a representative bin for NSHQ14, and was one of the most abundant bins in that well (Figure 6; Fones et al., 2019). Both bins also contained the acetoclastic methanogenesis pathway, as well as a nearly complete pathway for methanogenesis from methoxylated aromatic compounds, except for the gene for the first enzyme in the pathway, *mtoB* (Figure 6, Supplementary Table 3). The *mtoB* gene was detected in 51 other bins; 80% of these bins displayed the greatest abundance in the hyperalkaline or contact wells (Supplementary Table 3). A complete 2-methylcitrate cycle for propanoate degradation was detected in four bins, including a representative bin of WAB71 (a member of the Microtrichales) and a representative bin of WAB105 (*Acinetobacter junii*) (Figure 6).

### Porphyrins, Pigments, and Quinones

Fourteen features in the mass spec data were annotated as intermediates of pigment, porphyrin, and quinone biosynthesis pathways (Figure 5). A feature putatively annotated as protoporphyrin IX, a central metabolite in the production of heme and chlorophyll, was detected in all wells and was particularly abundant (3-5 orders of magnitude greater) in NSHQ14 (Figure 5). Another feature, putatively annotated as coproporphyrinogen III (a precursor to protoporphyrin IX) was detected in every well but displayed the lowest abundance in NSHQ14, by more than two orders of magnitude (Figure 5). Complete pathways for the biosynthesis of heme from glutamate and/or glutamyl-tRNA were annotated in bins detected in every well; however, bins containing complete or nearly complete (≥90%) heme pathways were generally more abundant in the hyperalkaline and contact wells (Figure 6).

Three features annotated as intermediates in ubiquinol (4-hydroxybenzoate) and menaquinone synthesis (2-succinyl-5-enolpyruvyl-6-hydroxy-3-cyclohexene-1-carboxylate and (1R,6R)-6-hydroxy-2-succinylcyclohexa-2,4-diene-1-carboxylate, intermediates in the synthesis of menaquinol and phylloquinol from isochorismate) were detected in all of the wells but were most abundant in NSHQ14 (Figure 5). Genes for the enzymes that produce these intermediates were detected in 14, 11, and six bins, respectively; these bins tended to be most abundant in the hyperalkaline or contact wells (Supplementary Table 3). The complete pathway for ubiquinone biosynthesis was detected in thirteen bins, including two representative bins of NSHQ14 (*Methanobacterium* and *Pseudomonas*) and two representative bins of WAB104 (a member of the Acidobacteriae, and *Pseudomonas alcaliphila*) (Figure 6). The complete pathway for menaquinone biosynthesis was detected in eleven bins, including a representative bin of every well except NSHQ14 (although one representative bin of that well, a member of the Moorellia, had 90% of the pathway) (Figure 6). The most commonly found cytochrome genes in the bins encoded for the cytochrome bd complex and NADH-quinone oxidoreductase. 41 of the 128 bins contained ≥90% of the genes encoding one or both of these cytochromes (Figure 6).

Features putatively annotated as (2’S)-deoxymyxol 2’-alpha-L-fucoside and glycosyl-4,4’-diaponeurosporenoate, metabolites produced in the biosynthesis of the carotenoids myxol 2’-dimethyl-fucoside and staphyloxanthin, respectively, were also detected in all the wells (Figure 5). The biosynthesis of (2’S)-deoxymyxol 2’-alpha-L-fucoside is a multi-step reaction that has not been fully characterized, but we were able to annotate the enzyme that catalyzes the conversion of (2’S)-deoxymyxol 2’-alpha-L-fucoside to myxol 2’-dimethyl-fucoside, beta-carotene hydroxylase, in 18 bins (Figure 6). Genes in the pathway for staphyloxanthin biosynthesis were annotated in nearly every bin, but the complete pathway was only detected in one bin, a member of the Anaerolineales that was most abundant in WAB71 (Figure 6).

A feature found exclusively in NSHQ14 and designated as a representative feature of that well was putatively annotated as tetraprenyl-beta-curcumene, an intermediate in the biosynthesis of a susquarterpene found in *Bacillus spp*. (Figure 5). The gene for tetraprenyl-beta-curcumene synthase was detected in seven bins, two of which were most abundant in WAB71, and the rest of which were most abundant in WAB105 (Figure 6).

Two features that were among the most abundant features in the mass spec data were annotated as intermediates of chlorophyll metabolism: magnesium protophorin and 13(1)-hydroxy-magnesium-protoporphyrin IX 13-monomethyl ester (Figure 5). Other putatively annotated features associated with photosynthesis and chlorophyll metabolism include phylloquinone, phytyl disphosphate, and phylloquinol (Figure 5). A feature annotated as either 3,4-dihydroxy-L-phenylalanine or N,N-dihydroxy-L-phenylalanine, precursors to pigments/cytochromes in fungi and plants, was also detected in all of the wells and was a representative feature of the alkaline/contact wells (Figure 5). Most of these features were relatively more abundant in the alkaline and contact wells. However, the feature putatively annotated as phylloquinone was exclusively detected in the hyperalkaline wells and was particularly abundant in NSHQ14 (Figure 5). Because the pathway for phylloquinol/phylloquinone synthesis and menoquinol synthesis involve the same/highly homologous enzymes, we could not distinguish between these pathways in the bins.

The complete pathway for chlorophyll biosynthesis was not detected in any bin (Supplementary Table 3). However, genes for two enzymes in this pathway, magnesium chelatase subunit I and anaerobic magnesium-protoporphyrin IX monomethyl ester cyclase, were annotated in 27 bins and 29 bins, respectively (Figure 6). Magnesium chelatase is a three-subunit enzyme responsible for inserting a Mg^+2^ into protoporphyrin IX in the first step of chlorophyll biosynthesis; genes for the other two subunits (H and D) were found in two and five bins, respectively, and none of the bins contained the genes for all three subunits of the enzyme (Supplementary Table 3). Anaerobic magnesium-protoporphyrin IX monomethyl ester cyclase produces 13(1)-hydroxy-magnesium-protoporphyrin IX 13-monomethyl ester in the chlorophyll biosynthesis pathway found in anaerobic photosynthetic bacteria. None of the bins contained the complete suite of genes required for photosynthesis, but two bins (one diagnostic) in the alkaline wells did contain the genes for an anoxygenic type-II reaction center (Supplementary Table 3). Thirteen bins contained the genes for the RuBisCo enzyme, nine of which also contained the complete Calvin-Benson-Bassham pathway for carbon fixation, but none of which included the anoxygenic type-II reaction center (Supplementary Table 3).

A feature putatively annotated as phytyl diphosphate, an intermediate of geranylgeranyl diphosphate biosynthesis, was also detected in all the wells. Geranylgeranyl diphosphate is a precursor to phylloquinol, chlorophyll, archaeol, and multiple pigments across all domains of life.

### Fatty Acids and Sterols

Eighteen features were annotated as fatty acids and intermediates in fatty acid pathways, five of which were designated characteristic features of the alkaline and contact wells by NMF analysis. These included four peaks annotated as dolichyl diphosphate, lipids involved in N-glycan production in archaea, which were detected in all the wells but were most abundant in NHSQ14 and WAB71. We also detected a feature putatively annotated as 12-oxo-9(Z)-dodecenoic acid (an intermediate of alpha-linolenic acid metabolism), a feature putatively annotated as 30-oxolanosterol (an intermediate of sterol biosynthesis in methylotrophs), and three features annotated as dodecanoic acid (Figure 5). There were also three features found in all the wells but particularly abundant in WAB71, which were annotated as prostaglandins by mummichog. MISA further confirmed the annotation of the latter two features as prostaglandin H2 or prostaglandin D2 (Figure 5; Supplementary Figure 1). However, we suspect that this annotation is incorrect and that these features are more likely intermediates or secondary metabolites of arachidonic acid and/or eicosapentanoic acid (see Discussion).

Genes for polyunsaturated fatty acid (PUFA) synthases (arachidonate and/or eicosapentaenoate synthase) were annotated in 18 of the bins analyzed in this study; however, a complete PUFA synthase operon (*pfaABCD*) was only found in two bins, one taxonomically annotated as belonging to the genus *Bellilinea* and the other as genus *Levilinea*, which are in the family Anaerolineaceae (phylum Chloroflexota, class Anaerolineae, order Anaerolineales) (Figure 6; Supplementary Table 3). The *pfaE* gene that encodes the phosphopantetheinyl transferase required for catalytic activation of acyl carrier protein domains was not present in these bins, but was found in two other bins belonging to phylum Chloroflexota, class Dehalococcoidia (Supplementary Table 3). Genes for fatty acid synthesis initiation and elongation (*fabB, fabD, fabF, fabG, fabH, fabI, fabZ*) and long chain acyl-CoA biosynthesis (*ACSL, fadD*), which produce the precursors to arachidonate and eicosapentanoate, were annotated in all but two of the representative bins (Figure 6).

### Amino Acid and Polyamine Metabolism

Thirteen features were putatively annotated as intermediates of amino acid/polyamine biosynthesis or degradation pathways (Figure 5). Most of these features were detected in all of the wells, with a few exceptions. A feature annotated as agmatine, an intermediate of arginine degradation and putrescine/spermidine biosynthesis, was present in every well except NSHQ14, and was designated a representative feature of the alkaline/contact wells by NMF analysis (Figure 5). Complete putrescine biosynthesis pathways were annotated in 12 bins; interestingly, nearly half of these bins displayed the greatest abundance in NSHQ14 (Figure 6). Conversely, a feature annotated as 5’-S-methyl-5’-thioinosine, a metabolite produced by the deamination of 5ʹ-methylthioadenosine (an intermediate of the methionine salvage cycle and spermidine biosynthesis), was detected in all the wells but was particularly abundant in NHSQ14 (Figure 5). The gene for enzyme that performs the deamination reaction, 5-methylthioadenosine/S-adenosylhomocysteine deaminase, was detected in 50 of the 128 bins analyzed in this study, including four representative bins of NSHQ14, one representative bin of WAB71, three representative bins of WAB55, two representative bins of WAB104, and one representative bin of WAB105 (Figure 6). This enzyme has also been implicated in the recycling of 5ʹ-deoxyadenosine and the biosynthesis of aromatic amino acids in methanogens (Miller et al., 2014). The methionine salvage cycle is also linked to the production of putrescine and spermidine. Only four bins contained ≥90% of the methionine salvage pathway, but 60 bins contained the portion of the pathway that results in 5ʹ-methylthioadenosine/spermidine biosynthesis (Figure 6).

Two representative features of NSHQ14 were putatively annotated as isethionic acid, a product of sulfoacetaldehyde degradation. These features were found only in NSHQ14 and WAB105, but at 8 orders of magnitude greater abundance in the former versus the latter well (Figure 5). Sulfoacetaldehyde is both a product of taurine degradation and an intermediate of coenzyme M biosynthesis. The gene for NADH-dependent sulfoacetaldehyde reductase, the enzyme that catalyzes the conversion of sulfoacetaldehyde to isethionic acid, was detected in 77 bins, including four representative bins from NSHQ14, one from WAB71, five from WAB55, three from WAB104, and two from WAB105 (Figure 6).

Two features annotated as either shikimate, 2-isopropylmaleate, or (2S)-2-isopropyl-3-oxosuccinate were detected in all of the wells but were most abundant in the alkaline and contact wells (Figure 5). Shikimate is an intermediate of the shikimate pathway for the production of chorismate, a precursor to aromatic amino acid and menaquinone/ubiquinone biosythesis. 2-Isopropylmaleate and (2S)-2-Isopropyl-3-oxosuccinate are intermediates of leucine biosynthesis. Two other features, annotated as alpha-isopropylmalate/2-isopropylmaleate/3-isopropylmalate, all of which are intermediates in leucine biosynthesis, were also detected in all the wells but most abundant in the alkaline wells (Figure 5). Complete shikimate and leucine biosynthesis pathways were detected in bins in every well (Figure 6). Nine of the 15 representative bins of the contact and alkaline wells contained one or both of these pathways, but only one of the eight representative hyperalkaline bins contained these pathways (Figure 6).

Other putatively annotated features related to amino acid metabolism include intermediates of lysine biosynthesis (2,3,4,5-tetrahydrodipicolinate), tryptophan degradation (3-hydroxyanthranilate and 2-aminomuconate), and phenylalanine metabolism (phenylacetic acid and 3-(3-hydroxyphenyl)propanoic acid) (Figure 5). Lysine biosynthesis pathways were detected in multiple bins (Figure 6). The complete pathway for tryptophan degradation via 3-hydroxyanthranilate to 2-aminomuconate was annotated in only one bin, which was taxonomically annotated as a member of the Bipolaricaulia (Supplementary Table 3). This bin was more abundant in WAB105 than any other well, but had a low relative abundance compared to other bins in that well. The gene *amiE*, which encodes the amidase enzyme that converts 2-phenylacetamide to phenylacetic acid, was detected in almost every bin; the gene *katG*, which encodes the catalase-peroxidase enzyme that converts phenylalanine to 2-phenylacetamide was detected in 43 bins (Figure 6). The enzyme in phenylalanine metabolism that produces 3-(3-hydroxyphenyl)propanoic acid has not been characterized, but the gene for 3-(3-hydroxy-phenyl)propionate hydroxylase, which converts 3-(3-hydroxy-phenyl)propionate acid to 2,3-dihydroxyphenylpropanoate in the same pathway, was detected in seven bins, only one of which was designated a representative bin of a well (WAB105) (Figure 6).

### Miscellaneous

A feature putatively annotated as pimelate, a precursor of biotin, was detected in every well, with a slightly greater relative abundance in the alkaline wells (Figure 5). The complete pathway for biotin synthesis from pimelate was annotated in only one bin, a member of the Thermodesulfovibrionales that displayed the greatest relative abundance in WAB71 (Figure 6). Twenty-one bins possessed four out of the five genes necessary for the pathway but were missing the gene for the enzyme 6-carboxyhexanoate-CoA ligase (*bioW*), which initiates the pathway by converting pimelate to pimeloyl-CoA (Supplementary Table 3). Two other bins contained this gene but otherwise lacked the complete pathway (Supplementary Table 3). The mechanism for the biosynthesis of pimelate is currently unknown, though it is speculated to begin with malonyl-CoA (Manandhar and Cronan, 2017). This same study found that deletion of the *bioW* gene results in a biotin auxotrophic phenotype in *Bacillus subtilis*.

Another feature, putatively annotated as 7-carboxy-7-carbaguanine (the precursor to preQ0 in preQ0 biosynthesis), was also detected in every well but was most abundant in WAB55 and WAB105 (Figure 5). A total of 36 bins contained the complete pathway for preQ0 biosynthesis, including one representative bin for NSHQ14, four representative bins for WAB55, two representative bins for WAB104, and three representative bins for WAB105 (Figure 6).

### Representative and Abundant Bins

Bins we identified as representative of alkaline wells belonged to the Bacillales (family Bacillaceae), Rhizobiales (family Rhizobiaceae), Pseudomonadales (*Acetinobacter junii* and *Pseudomonas alcaliphila*), Acidobacteriae, Bacteroidota (UBA10030), and Chloroflexota A (Figure 6). The most abundant bins in the alkaline wells also included members of the Thaumarchaeota, Actinobacteriota, Methylomirabilota, and Saccharimonadales (Figure 6). representative hyperalkaline bins included members of the Thermodesulfovibrionales, Bipolaricaulia (formerly candidate phylum OP1), Chloroflexota (Dehalococcoidia), Moorellia, Methanobacteriales, Microtrichales, Gemmatimonadetes, and Pseudomonadales (*Pseudomonas aeruginosa*) (Figure 6). Members of the Deinocococales, Burkholderiales, and Nitrospirales were also among the most abundant bins in the hyperalkaline wells (Figure 6). Bins representative of WAB55 (the only contact well sampled in this study) included Nitrospirales, Brocadiales, Chloroflexota (Anaerolineales), Gammaproteobacteria (UBA4486), Firmicutes (CSP1-3), and Poribacteria (Figure 6).

## Discussion

Previous studies have demonstrated clear trends correlating microbial diversity and metabolic capability to rock/fluid type in the Samail Ophiolite (Rempfert et al., 2017; Fones et al., 2019; Kraus et al., 2021; Fones et al., 2021). The chemistry of the subsurface fluids reflects both hydrologic context (i.e. the extent of equilibration with the atmosphere or host rock, or potential for mixing with other fluids) and the geology of the host rock (Rempfert et al., 2017; Leong et al., 2021). Multitracer experiments indicate that the higher pH (>9.3) fluids are older than their less alkaline counterparts, and their chemistry reflects a greater extent of serpentinization in the host rock (Paukert Vankeuren et al., 2019). Microbial community composition correlates with fluid geochemistry, with gabbro and alkaline peridotite fluids displaying a greater richness of microorganisms than the low-diversity communities found in hyperalkaline peridotite (Rempfert et al., 2017). Due to the covariance of multiple geochemical parameters, it is difficult to determine which of these parameters drives these correlations (Rempfert et al., 2017).

The extracellular mixture of compounds that constitutes the dissolved organic matter pool contains growth factors, substrates, and chemical signals that represent the fundamental relationship between microbes and their environment (Kujawinski, 2011; Douglas, 2020). Most primary metabolites tend to be retained within cells due to their charge at physiological pH values (Bar-Even et al., 2011). These metabolites are usually involved in multiple metabolic pathways and are common across clades of organisms (Peregrín-Alvarez et al., 2009). Secondary metabolites, however, are often more species-specific, and more likely to diffuse into the surrounding environment (Breitling et al., 2013; Covington et al., 2016). Secreted metabolites also act as a chemical language for microorganisms, whose transmission is in constant flux due to evolutionary and environmental pressures (Soldatou et al., 2019; Douglas 2020). The secretion of metabolites into the surrounding environment is a clear reflection of microbial activity, as the production of these compounds relies on the functional operation of metabolic pathways (Kell et al., 2005). Exometabolomics, or “metabolomic footprinting” (Kell et al., 2005), thus allows for the direct characterization of the molecular interaction between microbes and their environment (Sogin et al., 2019, Douglas, 2020). Metabolomics techniques, in conjunction with metagenomic data, have previously been used to describe the pool of intracellular and extracellular metabolites produced by microbial communities in the Coast Range Ophiolite Microbial Observatory (CROMO), a series of wells that provide access to a serpenization-influenced aquifer in the Coast Range of northern California (Seyler et al., 2020a). Here, we present a broad snapshot of the DOM pool in the Samail Ophiolite groundwater and identify some of the secondary metabolites produced by the microbial communities found there. The annotation of these features was supported by comparing putative compounds to the genes identified in the metagenomic bins.

Given that both microbial community structure and protein-coding gene abundance correlate with rock and fluid type (Rempfert et al., 2017; Fones et al., 2019; Kraus et al., 2021; Fones et al., 2021), we hypothesized that DOM composition would likewise correlate to these characteristics. Principal components analysis (Figure 1), NMF analysis (Figure 2), and procrustes analysis (Figure 4) comparing mass spec data across five wells lend support to this hypothesis. The DOM pools of NSHQ14, WAB71, and WAB55 are each distinguished from the other wells in these analyses, while the two alkaline wells analyzed (WAB104 and WAB105) contain DOM pools that are very similar to each other. TOC was highest in the hyperalkaline wells, lower in the alkaline wells, and lowest in the contact wells (Table 1). Correlating TOC to cell count resulted in an R^2^ value of 0.503, indicating that TOC is only partially driven by the amount of biomass present.

We hypothesize that the relative shortage of electron acceptors in the hyperalkaline wells leads to dissolved organics being degraded and remineralized more slowly in these environments (Fones et al., 2019). The higher TOC values in the hyperalkaline wells also may reflect the abiotic production of formate and/or small molecular weight organic compounds associated with serpentinization (Proskurowski et al., 2008; McCollom, 2013). For example, the features annotated as butanoic acid, which displayed the greatest abundance in the hyperalkaline wells, may be the result of abiotic organic synthesis (McCollom and Seewald, 2006), as we did not detect the pathway for butanoic acid production via fermentation in the metagenomic data. Similarly, relatively high abundances of features annotated as intermediates of propanoate degradation in the hyperalkaline wells raise the intriguing possibility that the abiotic production of propanoate could serve as a carbon source for the microbial communities found there (Shock and Schulte, 1998). Recent carbon analysis of serpentinized rocks obtained in the Oman Drilling Project shows that non-carbonate-carbon loadings are approximately 140 ppm to 360 ppm (Ternieten et al., 2021). Thus, there may be multiple primary sources of organic carbon in this system: autotrophic carbon fixation and turnover of biomass, heterotrophic utilization of reduced carbon accumulated in the rock during prior water/rock interaction, or ongoing abiotic organic synthesis.

Metagenomic data from the hyperalkaline wells indicates an enrichment of functional genes involved in C1 metabolisms such as CO utilization, methanogenesis, and acetogenesis (Fones et al., 2019; Colman et al., 2022). Our characterization of the DOC pool generally supports these findings. Features annotated as metabolites associated with methanogenesis tended to display greater relative abundance in the hyperalkaline wells, where both methane and methanogens have been detected by previous studies (Rempfert et al., 2017; Fones et al., 2019; Kraus et al., 2020). The features putatively annotated as intermediates of propanoate degradation (mentioned above) were also putatively annotated as cis-homoaconitate (Figure 5), an intermediate in the biosynthesis of coenzyme B, which reacts with coenzyme M to release methane during methananogenesis. The detection of isethionic acid in NSHQ14 may be linked to the degradation of sulfoacetaldehyde produced during coenzymeM biosynthesis, while the abundance of dolichyl diphosphate, an intermediate of N-glycan biosynthesis in archaea, may also be a result of the high abundance of methanogenic archaea in NSHQ14 (Fones et al., 2019; Fones et al., 2021).

The feature annotated as isocitrate was three orders of magnitude more abundant in the hyperalkaline wells than in the alkaline wells, and seven orders of magnitude higher in the hyperalkaline wells than in the contact well (Figure 5). Isocitrate has been implicated as a metabolic intermediate in some methanogens; for example, Methanosarcinales generates CO_2_ and 2-oxoglutarate from isocitrate via a partial oxidative TCA cycle (Anderson et al., 2009). This activity may serve as a source of CO_2_ for the production of methane by hydrogenotrophic methanogen such as *Methanobacterium*, which do not contain the pathway for 2-oxoglutarate generation via isocitrate (Anderson et al, 2009), but are abundant and active in NSHQ14 (Rempfert et al., 2017; Kraus et al., 2021). A bin annotated as *Methanobacterium* was the most abundant bin in NSHQ14, and designated as a representative bin of NSHQ14 by NMF analysis (Figure 6). This bin did not contain the gene for the NAD-dependent isocitrate dehydrogenase required for CO_2_ and 2-oxoglutarate generation via isocitrate, but 20 other bins did contain this gene, more than half of which were most abundant in the hyperalkaline wells (including one bin belonging to the Moorellia that was designated a representative bin of NSHQ14, and a likely homoacetogen) (Supplementary Table 3). A syntrophy relationship may exist between hydrogenotrophic methanogens and bacteria containing the partial oxidative TCA cycle, whereby the CO_2_ generated through this pathway fuels methanogenesis. Further studies, perhaps involving tracer/microcosm experiments, will be necessary to confirm if isocitrate generates CO_2_ for methanogenesis in the hyperalkaline wells.

Isocitrate is an important central metabolite in many processes, including glyoxylate and dicarboxylate metabolism, 2-oxocarboxylic metabolism, and the biosynthesis of signaling molecules and secondary metabolites. The glyoxylate cycle allows microorganisms to use C2 compounds such as acetate to satisfy cellular carbon requirements when simple sugars such as glucose are not available (Kornberg and Madsen, 1957; LaPorte et al., 1984). Microcosm experiments conducted in parallel to this study found that rates of assimilation/oxidation of acetate were higher than those of bicarbonate and carbon monoxide across all sampled wells, suggesting that acetate may be the preferred source of bioavailable carbon (Fones et al., 2019; Colman et al., 2022).

Metagenomic data have also shown an enrichment in genes involved in anaerobic metabolisms in hyperalkaline waters, while the alkaline wells were enriched in respiratory and electron transport chain genes (Fones et al., 2019). In contrast to the previously published metagenomic study, we found that features annotated as quinones and their precursors were generally enriched in NSHQ14, and to a lesser extent WAB71, compared to the contact and alkaline wells, with the exception of coproporphyrinogen III and phylloquinol (Figure 6). The abundance of these intermediates in the deeper wells may be indicative of slower turnover rates due to the extremophilic nature of this environment, or they may serve as intermediates of a process not yet characterized.

The abundance of features associated with plant and fungi pigments, including chlorophyll, in the alkaline and contact wells may reflect the greater influence of meteoric water in these wells (Nothaft et al., 2021). However, while the complete pathways for chlorophyll biosynthesis were not detected in the metagenomic data, the genes for the production of the intermediates we detected in the metabolome were found in multiple bins (Figure 6). In general, features related to the biosynthesis of pigments including carotenoids, quinones, and the chlorophyll intermediates may be linked to oxidative stress. Increased production of carotenoids, for example, has been observed previously in both neutrophilic (Lan et al., 2009) and extremophilic microorganisms (Tian and Hua, 2010; Mendelli et al., 2012) under oxidative stress conditions.

Of particular interest is the detection of several features putatively annotated as prostaglandins. These features were detected in all the wells, but were two to six orders of magnitude more abundant in WAB71 than in the other wells. We immediately questioned the validity of this annotation, as prostaglandins are signaling molecules found in higher eukaryotes. MISA analysis suggested that these features were either prostaglandin H_2_ or prostaglandin D2, both of which are involved in the human inflammatory response. However, prostaglandin H_2_ has a half-life of 90-100 seconds at room temperature (Human Metabolome Database), while prostaglandin D2 is found chiefly in brain and mast cells. These factors make it unlikely that these features are the result of contamination of our samples via casual contact. They may be prostaglandins produced by plants, but their abundance in WAB71, one of the most alkaline and least surface-influenced of the wells, makes this likewise unlikely.

We propose that these features may instead be secondary metabolites of arachidonic (ARA) or eicosapentaenoic acid (EPA). These polyunsaturated fatty acids serve as precursors to prostaglandins in higher eukaryotes, but are also found in some (mostly marine) bacteria belonging to the Gammaproteobacteria, Bacteroidetes, and Cyanobacteria. Eicosapentaenoic acid was annotated in the mass spec data during analysis with Progenesis QI, but MISA did not confirm this annotation (data not shown). Bacterial biosynthetic mechanisms of long-chain polyunsaturated fatty acids (LC-PUFAs) are not well understood and the functions of these lipids in bacteria have not been experimentally confirmed (Yoshida et al., 2016). It has been suggested that the primary function of LC-PUFAs is the adjustment of cell membrane fluidity under conditions of low temperature and/or high pressure (Kato and Nogi, 2001). EPA is required for growth at low temperatures and high pressures by some *Shewanella* species (Wang et al., 2004; Kawamoto et al., 2009; Usui et al., 2012), but a study of an EPA-deficient strain of *Photobacterium profundum* showed that monounsaturated or branched chain fatty acids could compensate for the loss of EPA (Allen et al., 1999). EPA and other LC-PUFAs have also been implicated in resistance to reactive oxygen species and oxidative stress (Nishida et al., 2006a; 2006b; 2007; 2010; Okuyama et al., 2008; Tilay and Annapure, 2012).

Four of the bins analyzed in this study contained ≥75% of the protein-coding genes required to produce arachidonate synthase and/or eicosapentaenoate synthase (Figure 6). Three of these bins were more abundant in WAB71 than any other well, and included two members of the Chloroflexota, order Anaerolineales (one of which was annotated *Bellilinea sp*.), and one member of the Desulfobacterota, family Desulfomonilaceae. The other bin was most abundant in WAB105, and was also a member of the Anaerolineales, *Livilinea sp*. The gene organization, order of genes, and composition of enzymatic domains in *pfa* gene clusters that produce LC-PUFAs appear to be highly conserved (Okuyama et al., 2007). *Pfa* gene homologs have also been identified in the genomes of 86 bacterial species belonging to 10 phyla, the products of which are largely unknown (Shulse and Allen, 2011). These homologs can be classified into 20 different gene types, which show a significant correlation with the ecological and physiological characteristics of the bacteria possessing the genes, suggesting that characterizing these secondary lipids may provide valuable insight into the ecological adaptation and evolution of microorganisms (Shulse and Allen, 2011). Lipid biomarkers have previously been characterized in rock samples obtained from the Samail Ophiolite; however, this study did not screen for LC-PUFAs (Newman et al., 2020). The *pfa* genes we detected in these bins may be homologs that produce an as-of-yet uncharacterized fatty acid product (Shulse and Allen, 2011).

## Conclusions

Characterization of the DOM pool in the Samail Ophiolite groundwater reveals the presence of numerous metabolic products and intermediates that may provide clues to the survival strategies of microbial communities in environments associated with serpentinization. We detected multiple metabolites linked to methanogenesis, including intermediates of coenzyme B and coenzyme M, byproducts of the degradation of these intermediates, and possible carbon sources for methane production, which were generally concentrated in the hyperalkaline wells. These results are encouraging, given the wealth of evidence as to the importance of methanogenesis in this environment (e.g., Rempfert et al., 2017; Fones et al., 2019; Kraus et al., 2021; Fones et al., 2021; Nothaft et al., 2021). Intermediates of the degradation of polysaccharides and amino acids were also detected and were particularly abundant in the hyperalkaline wells. We also found numerous features in the metabolomic data that suggest the importance of strategies for dealing with oxidative stress, such as pigments and PUFAs. Among the features we were able to annotate and confirm with metagenomic data, the pigments, porphyrins, fatty acids, sterols, and intermediates/byproducts of pathways associated with methanogenesis (such as coenzyme B and coenzyme M biosynthesis) were the compounds whose abundance was most strongly correlated to fluid type. In particular, pigments are a promising and intriguing biosignature of life in this system, due to both their specificity to particular processes and their longevity in the geological record (Schwieterman et al., 2015).

We chose methods for characterization that we hoped would allow us to capture as many DOM features as possible. These methods naturally have limitations. Many of the metabolic pathways of interest in these wells, in particular those involved with C1 metabolisms, consist of intermediate compounds that are involved in multiple overlapping pathways (Peregrín-Alvarez et al., 2009), many of which are too small to be detected using the liquid chromatography methods used in this study. These compounds are also cycled rapidly and retained within the cell (Bar-Even et al., 2011), precluding them from an exometabolome analysis such as this one. Additionally, many of the features we detected could not be annotated using available metabolite databases-a caveat which reflects both the under-characterization of microbial secondary metabolites on the whole, but also the uniqueness of the environment and the microbial communities that inhabit it.

Linking DOM composition to environmental characteristics has exciting implications for the search for potential biomarkers for life fueled by serpentinization, in particular because the greatest numbers of unique features were found in wells that are the most heavily influenced by serpentinization. Metabolomics studies of environments of astrobiological significance, such as serpentinizing rock, hold great promise in the search for biomarkers that may be used as evidence of life on other worlds (Seyler et al., 2020b). Further efforts to characterize the unique features detected in hyperalkaline fluids in the Samail Ophiolite may thus provide researchers with the means to detect life in serpentinizing systems elsewhere in our Solar System, such as the subsurface of Mars or the icy moons of Jupiter and Saturn. Such techniques hold the possibility of uncovering biomarkers of processes and relationships that are representative of habitats influenced by serpentinization, aiding the search for life in similar environments on other worlds in our Solar System.

## Acknowledgements

The authors would like to thank Kaitlin Rempfert, Eric Ellison, Laura Beuter, and Eric Boyd for assistance with sample collection, Juerg Matter for help with equipment acquisition, permitting, and sample export, and the Ministry of Regional Municipalities and Water Resources in the Sultanate of Oman for allowing sampling and export of well waters. We also wish to thank A. Daniel Jones and Anthony Schilmiller for their assistance with performing QToF-LC/MS/MS, and Elizabeth Kujawinski for her guidance regarding data analysis and interpretation. This work was supported by a grant from the NASA Astrobiology Institute (NNA15BB02A) to AST, JSR, EB, and MOS, and a grant from the Deep Carbon Observatory - Deep Life Community (#2015-14084) to MOS.

---

**Supplementary Table 1**. All features identified as representative of the three well types (“levels”) determined by non-negative matrix factorization (NMF) analysis. Annotations acquired by mummichog are included where available. Features highlighted in bold were annotated by Progenesis QI using the ChemSpider Database (Supplemental Table 2).

**Supplementary Table 2**. Representative features that could be annotated by Progenesis QI using the ChemSpider database.

**Supplementary Table 3**. KOs for all genes annotated by KEGGDecoder in bins included in this study.

